# When does diet matter? The roles of larval and adult nutrition in regulating adult size traits

**DOI:** 10.1101/2019.12.18.880708

**Authors:** Gonçalo M. Poças, Alexander E. Crosbie, Christen K. Mirth

## Abstract

Adult body size is determined by the quality and quantity of nutrients available to animals. In insects, nutrition affects adult size primarily during the nymphal or larval stages. However, measures of adult size like body weight are likely to also change with adult nutrition. In this study, we sought to the roles of nutrition throughout the life cycle on adult body weight and the size of two appendages, the wing and the femur, in the fruit fly *Drosophila melanogaster*. We manipulated nutrition in two ways: by varying the protein to carbohydrate content of the diet, called macronutrient restriction, and by changing the caloric density of the diet, termed caloric restriction. We employed a fully factorial design to manipulate both the larval and adult diets for both diet types. We found that manipulating the larval diet had greater impacts on all measures of adult size. Further, macronutrient restriction was more detrimental to adult size than caloric restriction. For adult body weight, a rich adult diet mitigated the negative effects of poor larval nutrition for both types of diets. In contrast, small wing and femur size caused by poor larval diet could not be increased with the adult diet. Taken together, these results suggest that appendage size is fixed by the larval diet, while those related to body composition remain sensitive to adult diet. Further, our studies provide a foundation for understanding how the nutritional environment of juveniles affects how adults respond to diet.

## 1. INTRODUCTION

Diet is known to affect the body size of most animals, which in turn has lifelong effects on an animal’s reproductive capacity and fitness (Simpson and Raubenheimer, 2012). While body size is largely thought to be a product of growth in response to diet during the juvenile stages (Callier and Nijhout, 2013; Nijhout, 2003; Nijhout et al., 2014), at least some adult measures of size might change with the adult diet. Because of their rigid exoskeleton, the size of insect appendages is largely fixed by the time they reach metamorphosis. However, internal structures are not as rigidly constrained. For instance, the accumulation of carbohydrates and lipids in the fat body and hemolymph are dynamic and are affected by the adult diet (Skorupa et al., 2008; Verdú et al., 2010), and these changes in body composition are likely to affect adult weight. Similarly, ovary size is affected by the composition of the adult diet (Bong et al., 2014; Verdú et al., 2010). This raises the question, to what extent are measures of adult size fixed by the larval diet, and how much can these be modified by the composition of the adult diet?

The growth of adult structures, such as the reproductive organs and the wings, is determined by the amount of nutrition available in nymphal or larval stages (Araújo et al., 2012; Awmack and Leather, 2002; Boggs and Freeman, 2005; Chaudhury, 1989; Colasurdo et al., 2009; Helm et al., 2017; Nijhout, 2003; Shafiei et al., 2001). This is because these adult structures grow within the larval body, and are subject to the same circulating hormones that drive growth in the larval tissues and increase larval mass (Nijhout et al., 2014). Further, the adult structures of insects like beetles, butterflies, and moths often further depend on the nutritional conditions of the larval stages, as these dictate the amount of stored nutrients that will be available to sustain growth throughout metamorphosis (Emlen and Nijhout, 1999; Nijhout, 2003; Nijhout and Grunert, 2010). In this way, the quantity of food available to the larva dictates the final size of structures like the insect appendages.

While quantity of food is certainly important, the quality of the food also matters. Studies over the past two decades have illustrated that the nutrient composition of diets impacts a wide range of adult traits, including size related traits (Blackmore and Lord, 2000; da Silva Soares et al., 2017; Gobbi et al., 2013; Grangeteau et al., 2018; Kutz et al., 2019; Magnarelli et al., 1982; Matavelli et al., 2015; Rodrigues et al., 2015; Sentinella et al., 2013; Shingleton et al., 2017; Souza et al., 2019; Tomberlin et al., 2002). For herbivorous and omnivorous insects, the nutrients that contribute the most to energy gain include protein and carbohydrate, with lipid playing a minor role (Behmer, 2009; Lee et al., 2008). Predatory insects consume very little carbohydrate, and their primary sources of energy include protein and lipids (Jensen et al., 2012; Mayntz et al., 2005). A wide range of insects, from cockroaches, locusts, beetles, and flies, are known to adjust their feeding to acquire precise combinations of these macronutrients to optimize life history trait outcomes (Fanson and Taylor, 2012; Jensen et al., 2012; Jones and Raubenheimer, 2001; Lee et al., 2008; Mayntz et al., 2005; Simpson et al., 1991).

The balance of these macronutrients in the diet during the nymphal or larval stages can have an important impact on size-related traits. For most insects, size traits tend to correlate with the amount of protein available in the larval diet (da Silva Soares et al., 2017; Kutz et al., 2019; Matavelli et al., 2015; Rodrigues et al., 2015; Sentinella et al., 2013; Shingleton et al., 2017). This has lead authors to conclude that total nutrient availability, in other words total calories, is not the only thing that matters for body size.

Macronutrient balance in the adult diet also shapes a number of important life history traits. For instance, lifespan is increased on diets that have lower protein to carbohydrate (P:C) ratios, while egg production is maximized on higher P:C ratios (Carvalho et al., 2005; Fanson et al., 2012; Fanson and Taylor, 2012; Lee et al., 2008). Additionally, nutrition can induce changes in body composition. When adults are feed on high sugar, low yeast diets, they increase the triglyceride and decrease the protein content in their bodies (Skorupa et al., 2008). While these changes in body composition are likely to affect size measures like adult weight, the relative contribution of larval and adult diet to metrics of adult size has not been explicitly examined.

In our study, we tested the effects of diet across life stages on appendage size and body mass in adult *Drosophila melanogaster* fruit flies. In this insect, the caloric content and the concentration of protein and carbohydrate of the larval diet affects many adult size traits, such as body weight and appendage size (Bakker, 1959; David, 1970; Kutz et al., 2019; Rodrigues et al., 2015; Shingleton et al., 2017). To test the relative contribution of larval and adult diet to measures of adult size, we performed two experiments. In the first, we altered the macronutrient composition of the larval and the adult diets by altering the P:C ratios of the diet while maintaining constant caloric densities. In the second experiment, we altered the caloric density of the larval and adult diets, while maintaining the P:C ratios constant. These findings allow us to parse out the relative effects of larval and adult diet on our adult body size measures, and to assess which type of diet manipulation affects these measures the most. These studies provide insights into how dietary composition across life stages impacts adult size traits in insects.

## 2. METHODS

### 2.1. Fly Stock

For this study, we used the wild-type Oregon R strain of *Drosophila melanogaster*. Flies were maintained under constant temperature (25°C), humidity (60%) and a 12:12 light-dark cycle, and fed ad libitum with sucrose-yeast diet containing 10 g of agar, 100 g of yeast extract and 50 g of sucrose in 1000 ml of water. To prevent bacterial and fungal growth, we added 3% Nipagen and 0.3% (v/v) propionic acid to the cooled mixtures.

### 2.2. Experimental Diets

All experimental diets were based on manipulating the ingredients in the sucrose-yeast diet, which contains a protein to carbohydrate (P:C) ratio of 1:2 and a caloric concentration of 495.36 calories/L. We conducted two experiments where larvae and adults were offered diets of different quality: a macronutrient dilution experiment and a caloric dilution experiment. For all the diets, yeast extract was used as protein. Both sugar and yeast extract contain carbohydrate. All pre-weighed dry ingredients were dissolved in sterile distilled water and stirred for 5-10 min. To set the diet, we added 2 g of agar to the suspension before autoclaving it for 50 min. Nipagen and propionic acid solutions were added to the food to a final concentration (v/v) of 3% and 0.3%, respectively.

For the macronutrient restriction experiment, we generated variation in the macronutrient composition of the food by making one of two diets: the standard sucrose-yeast diet with a protein to carbohydrate (P:C) ratio of 1:2, and a protein-poor diet with a P:C ratio of 1:10. To alter the P:C ratio, we changed the balance of yeast and sucrose such that the 1:10 diet contained 25 g of yeast and 108.5 g of sucrose per litre of food (Table 1). The caloric density was held constant across both diets (495.36 calories/L).

**Table 1:** Composition of experimental diets.

| Control food (1:2 and 100%) |  |  |  |  |  |  |  |
| --- | --- | --- | --- | --- | --- | --- | --- |
|  | Protein per 100g | Carbs per 100g | Amount (g) in 1L | Protein in 1L | Carbs in 1L | P + C | Calories |
| Agar | 0 | 0 | 10 | 0 | 0 |  |  |
| Sugar | 0 | 100 | 50 | 0 | 50 |  |  |
| Yeast | 45 | 33 | 100 | 45 | 33 |  |  |
| Total |  |  |  | 45 | 83 | 128 | 495 |
| Macronutrient Restriction (1:10) |  |  |  |  |  |  |  |
| Agar | 0 | 0 | 10 | 0 | 0 |  |  |
| Sugar | 0 | 100 | 108.5 | 0 | 108.5 |  |  |
| Yeast | 45 | 33 | 25 | 11.25 | 8.25 |  |  |
| Total |  |  |  | 11.25 | 116.75 | 128 | 495 |
| Caloric Restriction (25%) |  |  |  |  |  |  |  |
| Agar | 0 | 0 | 10 | 0 | 0 |  |  |
| Sugar | 0 | 100 | 12.5 | 0 | 12.5 |  |  |
| Yeast | 45 | 33 | 25 | 11.25 | 8.25 |  |  |
| Total |  |  |  | 11.25 | 20.75 | 32 | 124 |

For the caloric restriction experiment, we created a low-calorie diet by diluting the sucrose-yeast diet with 1% agar to 25% the concentration of the standard diet, maintaining the P:C ratio constant at 1:2 (Table 1). Our standard sucrose-yeast diet was used as our high calorie diet for comparisons.

### 2.3. Rearing conditions

Approximately 100 flies (70 females and 30 males) were transferred into egg laying chambers (100 ml plastic cups) where they were left to lay eggs on apple juice plates (30% agar, 30% sucrose, and 40% apple juice in 60 mm Petri dishes) seeded with live yeast paste for 4 hours. From these dishes, we collected 40 eggs, washed them in PBS to remove any food or yeast paste, and transferred them to fly vials containing 7mL of one of two test diets for either the macronutrient dilution or caloric dilution experiments (Figure 1). The animals were allowed to develop through the three larval stages and undergo metamorphosis in the vials, before eclosing as adults at 25 °C in a climate-controlled room under 60–70% humidity and a 12:12 light-dark cycle.

**Figure 1.**
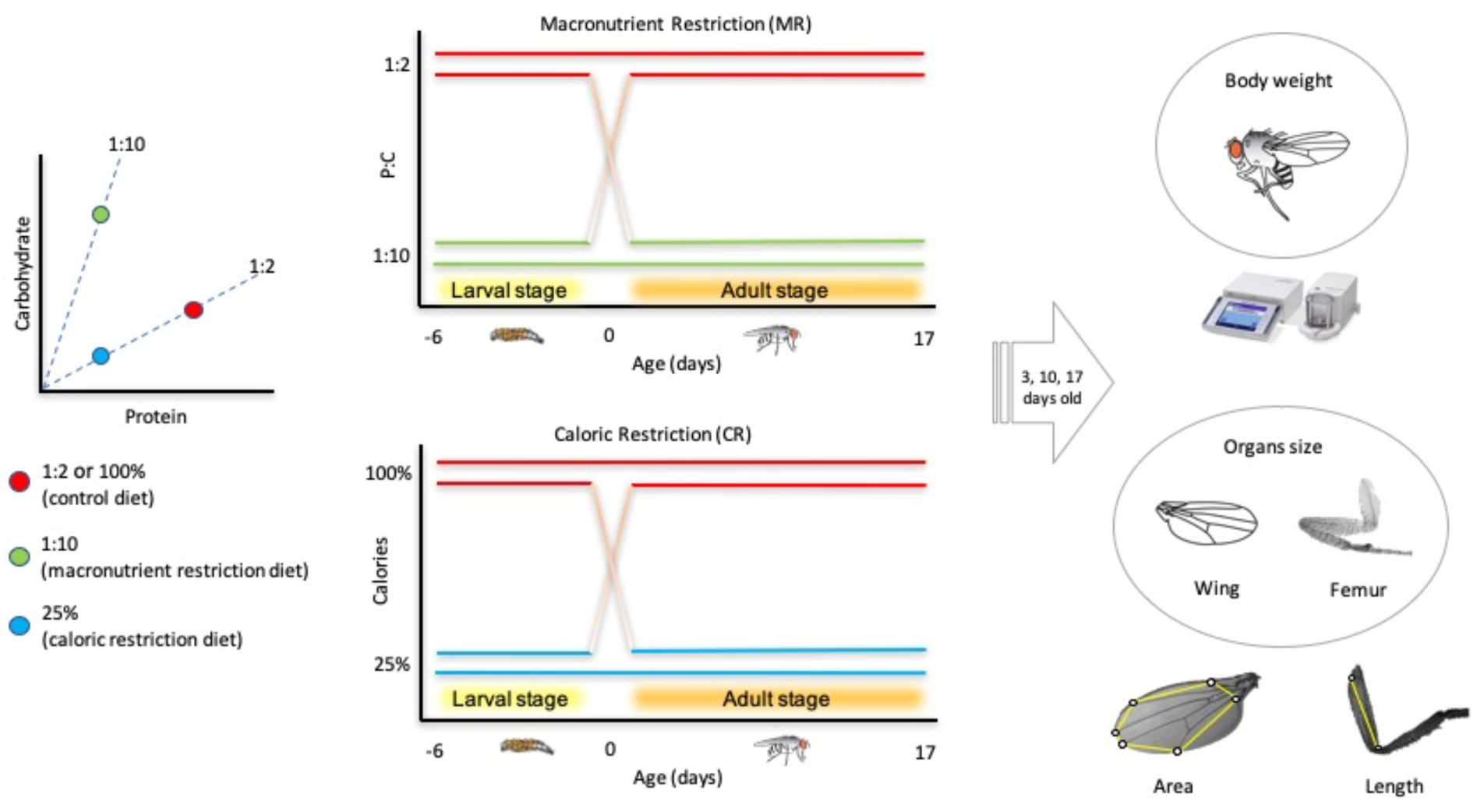
We performed two experiments to investigate the effect of diet on measurements of adult body size: a macronutrient restriction experiment and a caloric restriction experiment. The effect of diet on adult traits was evaluated by varying either the macronutrient composition or caloric density of diets available during the larval and adult stages. In the macronutrient dilution experiment, we offered animals either a control diet containing a P:C ratio of 1:2 and a caloric content of 495 calories/L (100%) or a low P:C diet also containing 495 calories/L but a P:C ratio of 1:10. For the caloric dilution experiment, we reared animals on either the control diet or on a low-calorie diet (25%), both of which had a P:C ratio of 1:2. In both cases larvae were offered one of two experimental diets. After eclosing, the adult flies (males and females) were either maintained on the same food or switched to the alternate diet. We maintained 30 adult flies per each vial (15 males and 15 females) and all individuals were maintained on their diets until sampling. Samples were weighed and dissected to determine body mass and organ size on days 3, 10, and 17 after eclosion.

After eclosing, 30 adult animals (15 males and 15 females) were transferred to a fresh vial containing either the same diet as that on which they were reared or the alternate diet (Figure 1). This meant that for each experiment there were four food treatment types: two treatments where the larvae were provided the same diet as adults and two treatments where the larval diet differed from the adult diet (Figure 1).

### 2.4. Measurements of adult body parts

We next explored whether macronutrient restriction or caloric restriction would differentially affect the adult body weight and the size of the adult appendages. We collected, weighed, and preserved adult animals in 80% ethanol from each of the vials 3, 10, and 17 days after they eclosed from the puparium. At each time point, we dissected 30 flies per treatment, 15 males and 15 females. We dissected the appendages off the bodies in SH solution (70% ethanol, 30% glycerol), using a Leica MZ75 scope to remove the right wing and right front leg. The appendages were mounted in 100% glycerol solution onto microscope slides. Digital images of the wings and femurs were captured using a Leica M165 FC stereoscope and a Leica DFC450 C digital camera.

All images were processed using ImageJ software. Wing area was estimated using landmarks from the veins (Figure 1) and femur length was measured using anterior and posterior landmarks (Figure 1).

### 2.5. Statistical analysis

All statistical analyses were conducted using RStudio (version 1.1.442). To analyse the effects of either macronutrient or caloric restriction on our measurements of adult size, we fit the data with generalized linear models using age, sex, and either P:C ratio of the larval and adult diets or caloric content of the larval and adult diets as fixed effects, including all possible interactions. To linearize the data, we applied a log10 transformation to all size measurements. All data and scripts are available on Figshare (DOI:10.26180/5df6ed2dc5ed5).

## 3. RESULTS

The aim of this study is to explore the relative roles of larval and adult diet on different measures of adult body size. We hypothesized that because adult body composition can change with adult diet (Galikova and Klepsatel, 2018), adult body weight would be sensitive to the composition of both the larval and adult diet. In contrast, we hypothesized that the size of sclerotized appendages would not be able to change in the adult, and thus would only be affected by the composition of the larval diet.

To test each hypothesis, we subjected animals to one of two dietary treatments. In the first, we subjected the larvae and adults to one of two macronutrient compositions, which included diets with P:C ratios of either 1:10 or 1:2. The second experiment exposed larvae and adults to diets varying in their caloric density, either 25% or 100% of the calories of standard sucrose/yeast medium. We then measured the effects of the larval and adult diets on male and female body weight, wing area, and femur length at three different adult ages: 3, 10 and 17 days old (Figure 1).

### 3.1. The effects of diet on adult body weight

#### 3.1.1. Macronutrient Manipulation

When we modified the macronutrient composition of the diet, we found that adult body weight varied with age, sex, larval diet, and adult diet. The largest amount of variance in adult weight was explained by sex, with females weighing more than males for all treatments (Figure 2A, Table 2). In addition, we found significant interactions between age and sex, larval diet and sex, and adult diet and sex (Table 2). Because both larval and adult diet appeared to effect adult weight differently in males and females, we analysed the data for each sex separately.

**Table 2:** The effect of dietary manipulations on adult body weight.

|  | Macronutrient Dilution |  | Caloric Dilution |
| --- | --- | --- | --- |
|  | Df | Chisq | Chisq |
| Age | 1 | 32.84 *** | 54.34 *** |
| Larval Diet | 1 | 588.03 *** | 43.36 *** |
| Adult Diet | 1 | 79.74 *** | 65.36 *** |
| Sex | 1 | 1847.44 *** | 1503.32 *** |
| Age : Larval Diet | 1 | 12.21 *** | 28.74 *** |
| Age : Adult Diet | 1 | 36.43 *** | 1.61 |
| Larval Diet : Adult Diet | 1 | 3.39 | 1.41 |
| Age : Sex | 1 | 13.17 *** | 2.024 |
| Larval Diet : Sex | 1 | 26.03 *** | 0.63 |
| Adult Diet : Sex | 1 | 25.78 *** | 40.23 *** |
| Age : Larval Diet : Adult Diet | 1 | 0.28 | 0.0026 |
| Age : Larval Diet : Sex | 1 | 2.99 | 1.37 |
| Age : Adult Diet : Sex | 1 | 1.45 | 0.48 |
| Larval Diet : Adult Diet : Sex | 1 | 0.55 | 0.12 |
| Age : Larval Diet : Adult Diet : Sex | 1 | 0.27 | 0.93 |

Significance codes: \* $p < 0.05$ , \*\* $p < 0.01$ , \*\*\* $p < 0.001$

**Figure 2.**
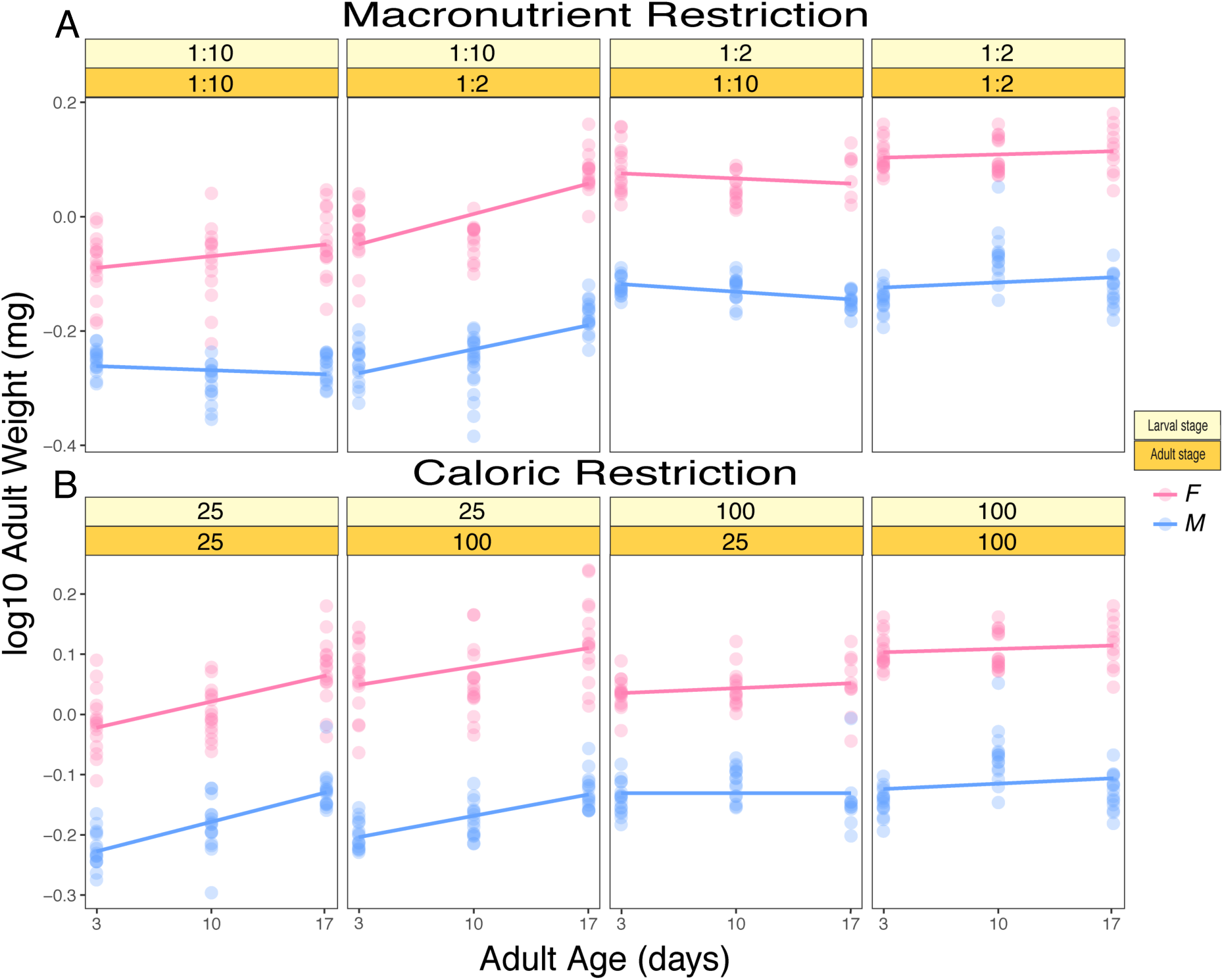
Adult body weight is affected by the macronutrient quality and caloric content of the diets available during the larval and adult stages. A) shows how adult body weight responds to the macronutrient composition of the larval and adult diets. B) shows how adult body weight responds to the caloric composition of the larval and adult diets. The stage at which treatments were imposed are indicated in the yellow-toned boxes: the larval nutrition in light yellow and the adult nutrition in darker yellow. The blue dots and lines represent the data for the adult males and the pink dots and lines represent the data for the adult females.

For both males and females, adult body weight was more affected by the P:C ratio of the larval diet than by the adult diet (Figure 2A, Table 3). In both cases, larvae that were fed on the control diet, which contained a P:C ratio of 1:2, had significantly higher adult body weights than those reared on the low P:C ratio (1:10) diet (Figure 2A, Table 3).

**Table 3:**
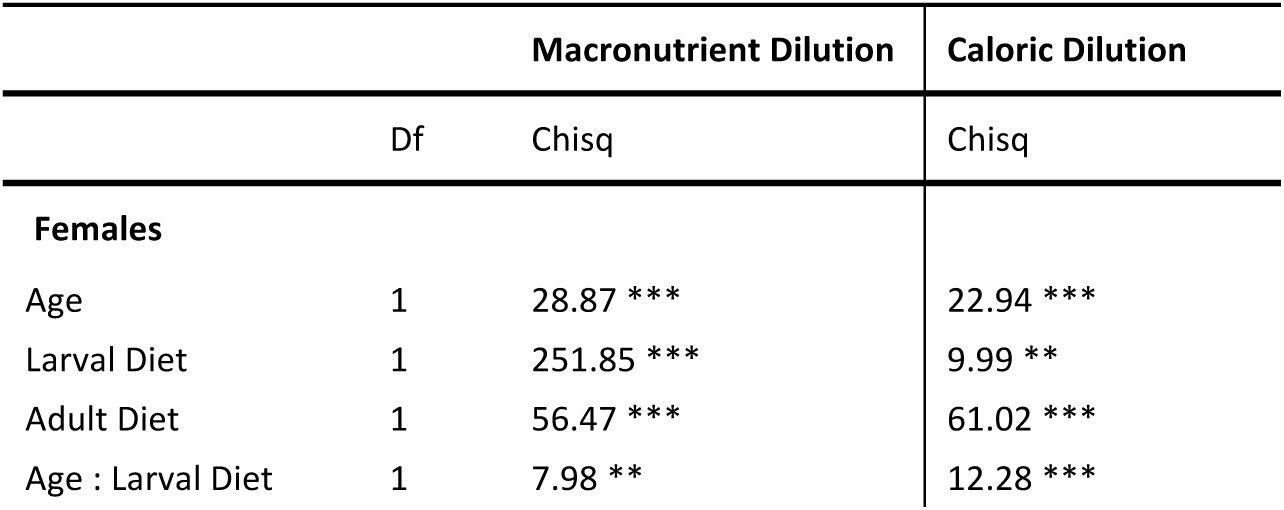
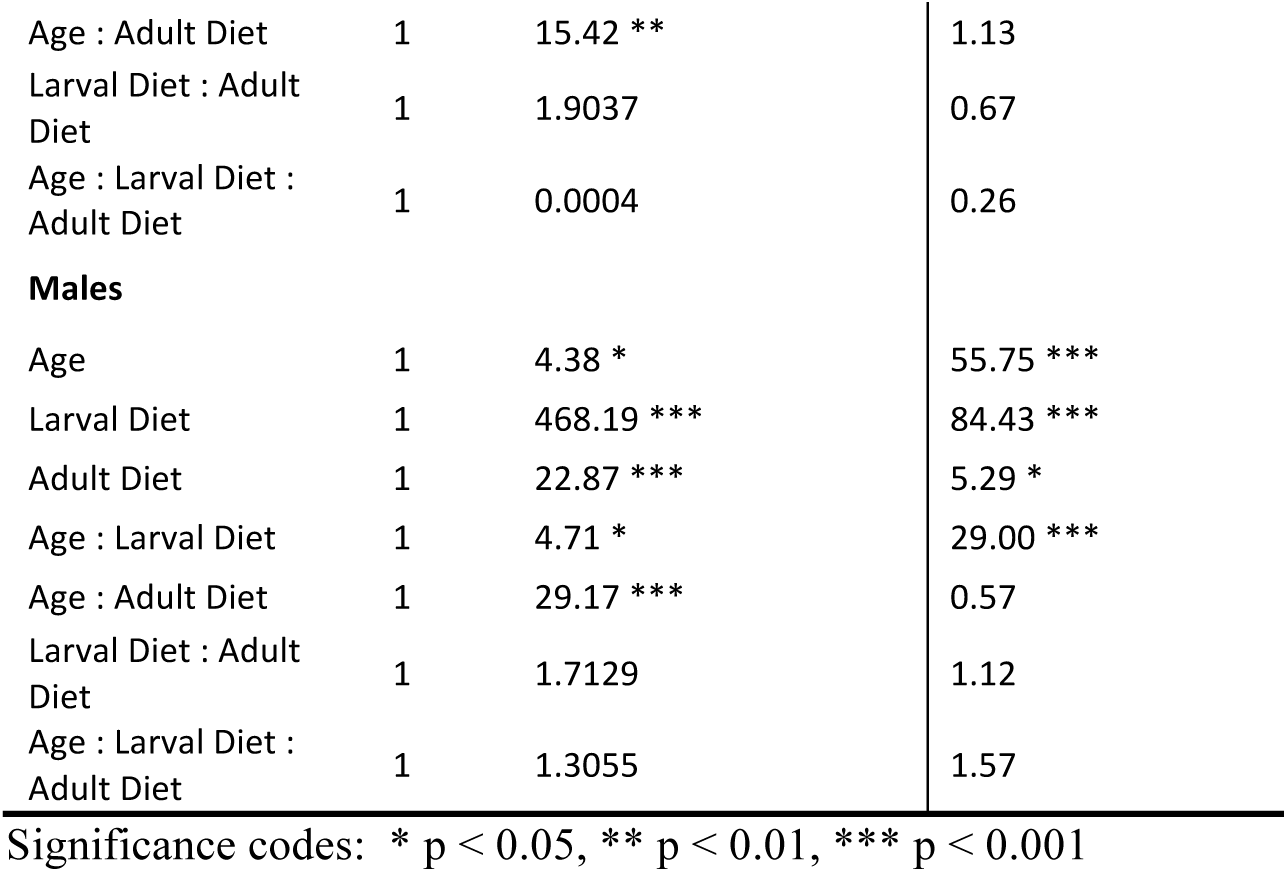
The effect of dietary manipulation on adult body weight in females and males.

While larval diet explained more of the variation in adult body weight, adult diet did affect adult body weight in both males and females. Males and females offered the 1:2 diet as adults were significantly heavier than those on the 1:10 diet (Figure 2A, Table 3).

Interestingly, the interactions between age and larval diet and age and adult diet were significant for both males and females. This was because, in both sexes, larvae that were reared on the 1:10 diet showed an increase in weight with age, whereas those reared on the 1:2 diet did not (Figure 2A, Table 3). Furthermore, for both sexes adult weight increased more with age for adults on the 1:2 diet than for adults on the 1:10 diet (Figure 2A, Table 3). The increase in adult body weight with age seemed to be less pronounced in males when compared to females, which would explain the significant interactions between sex and age (Table 2). The three-way interaction between age, larval diet, and adult diet was not significant in either sex, meaning that the larval diet did not influence how adult diet affected adult body weight with age.

#### 3.1.2. Caloric Manipulation

Next, we subjected both larvae and adults to diets that varied in their caloric density. Similar to our results for macronutrient manipulations, we found that sex explained the greatest amount of variation in adult body weight (Figure 2B, Table 2). Further, the significant interaction between adult diet and sex indicates that males and females differ in how their body weight responds to the adult diet. As such, we analysed the results for males and females separately.

In contrast to when we varied the macronutrient composition of the diet, we found that the caloric density of the adult diet explained more variation in female weight than that of larval diet (Figure 2B, Table 3). For males, however, larval diet explained more variation in body weight than adult diet (Figure 2B, Table 3). In both cases, body weight was significantly higher for males and females reared on the higher caloric (100%) food as larvae, and also higher for adults fed on the higher calorie diet (Figure 2B, Table 3).

Body weight increased significantly with age for both males and females, and larval diet significantly affected the extent of this response in both sexes, resulting in a significant interaction between age and the calorie content of the larval diet (Figure 2B, Table 3). The caloric density of the adult diet did not impact the increase in body weight with age.

In sum, our results from both the macronutrient manipulation and the caloric manipulation experiments support our hypothesis that both larval and adult diet contribute to variation in adult body weight.

### 3.2. The effects of dietary manipulations on appendage size: wing area and femur length

#### 3.2.1. Macronutrient Manipulation

When we manipulated the macronutrient content of the larval and adult diet, we found that the size of both appendages depended on the macronutrient quality of the diet available in the larval stages and on sex (Table 4). When larvae were reared on the higher protein diet (1:2), they had larger wings and femurs (Figure 3A & 4A, Table 4). For femur length, the age of the adult and the P:C ratio of the adult diet also affected size. This suggests that the femur does change size with age.

**Table 4:**
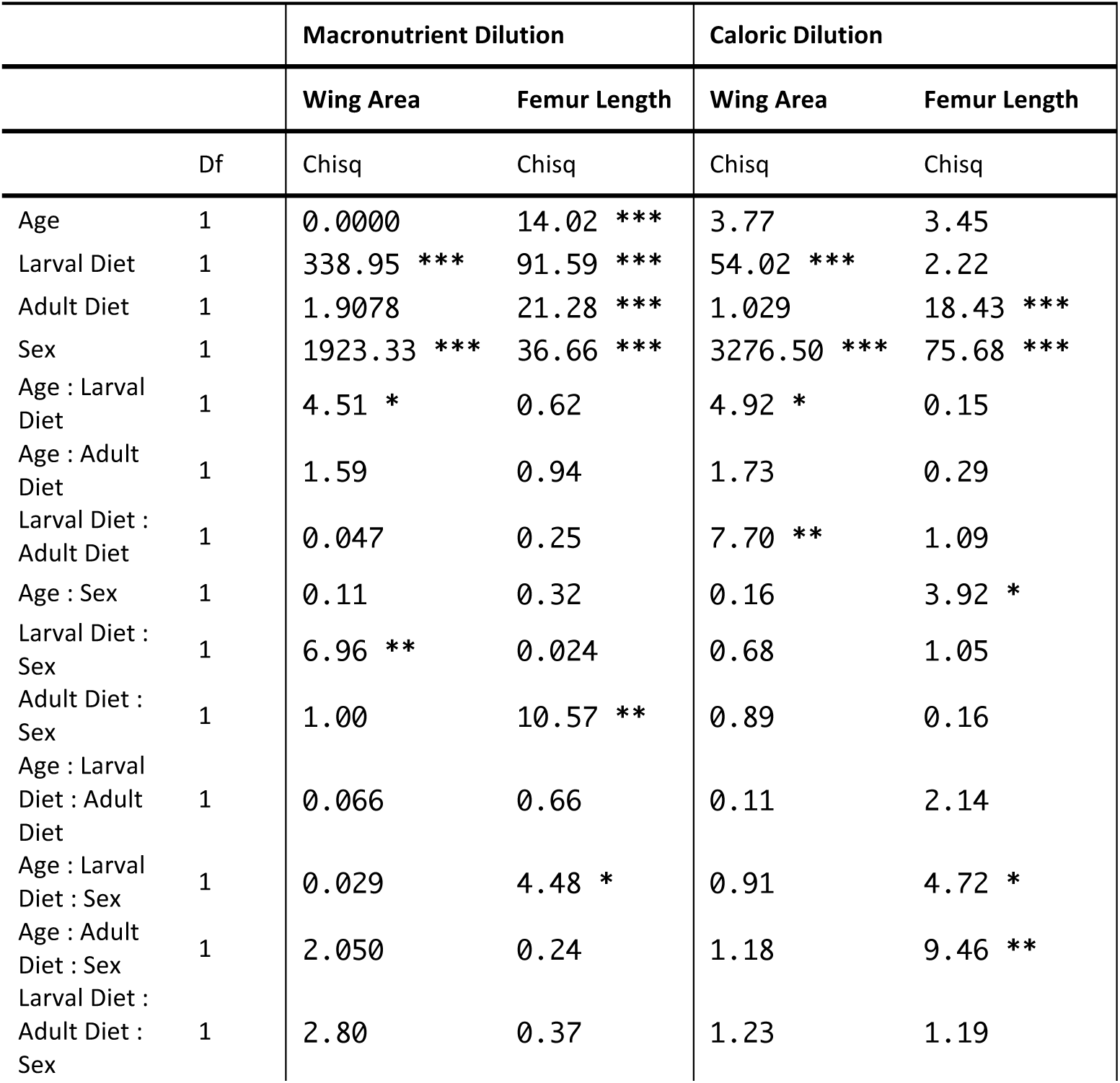
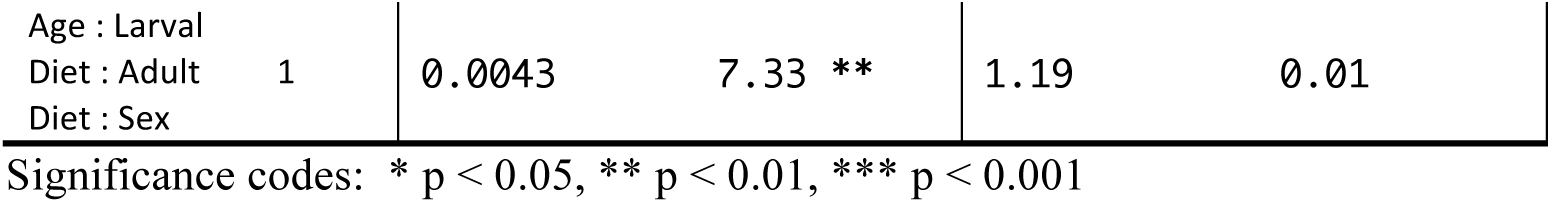
The effects of manipulating the larval and adult diets on wing area and femur length.

**Figure 3.**
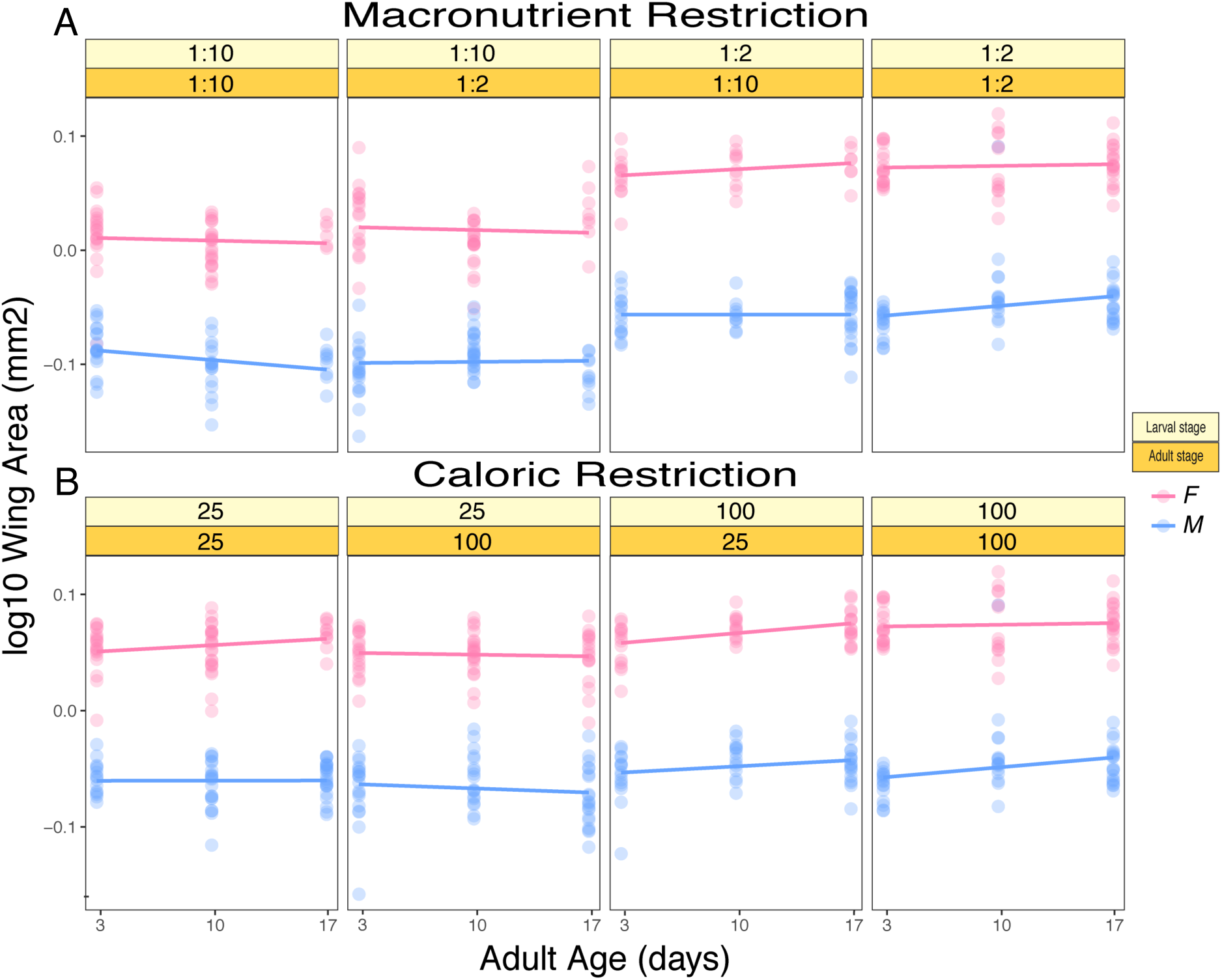
The size of the wings is mainly determined by the diet available during the larval stage. A) shows the effects of the macronutrient composition of the larval and adult diet on wing area. B) shows the effects of the caloric composition of the larval and adult diet on wing area. The stage at which treatments were imposed are indicated in the yellow-toned boxes: the larval nutrition in light yellow and the adult nutrition in darker yellow. The blue dots and lines represent the data for the adult males and the pink dots and lines represent the data for the adult females.

Sex contributed significantly to variation in appendage size for both wings and femurs, with females having larger appendages than males overall (Figure 3A & 4A, Table 4). For wings, we identified significant interactions between age and larval diet and larval diet and sex (Table 4). Femur size depended on interactions between the P:C ratio of the adult diet and sex, a three-way interaction between age, the P:C ratio of the larval diet, and sex, and a complex four-way interaction between age, the P:C ratio of the larval diet, the P:C ratio of the adult diet, and sex (Table 4). This would mean that the way that femurs change in size with age depends on both the larval and adult diet. To gain more insight into the nature of these effects, we analysed the data for each sex separately.

For females, the size of the wing was only significantly affected by the P:C ratio of the larval diet. Females fed as larvae on the protein-rich food, containing a P:C ratio of 1:2, had larger wings than those maintained as larvae on the lower P:C ratio (1:10) (Figure 3A, Table 5). Similarly, females reared on the more protein-rich diet had larger femurs than those reared on the lower P:C ratio food (1:10) (Figure 4A, Table 5). Interestingly, the P:C ratio of the adult diet also affected mean femur length in females, with females offered the higher protein diets showing larger femur sizes overall (Figure 4A, Table 5). Thus, while wing size did not change significantly with adult diet, femur size did.

**Table 5:**
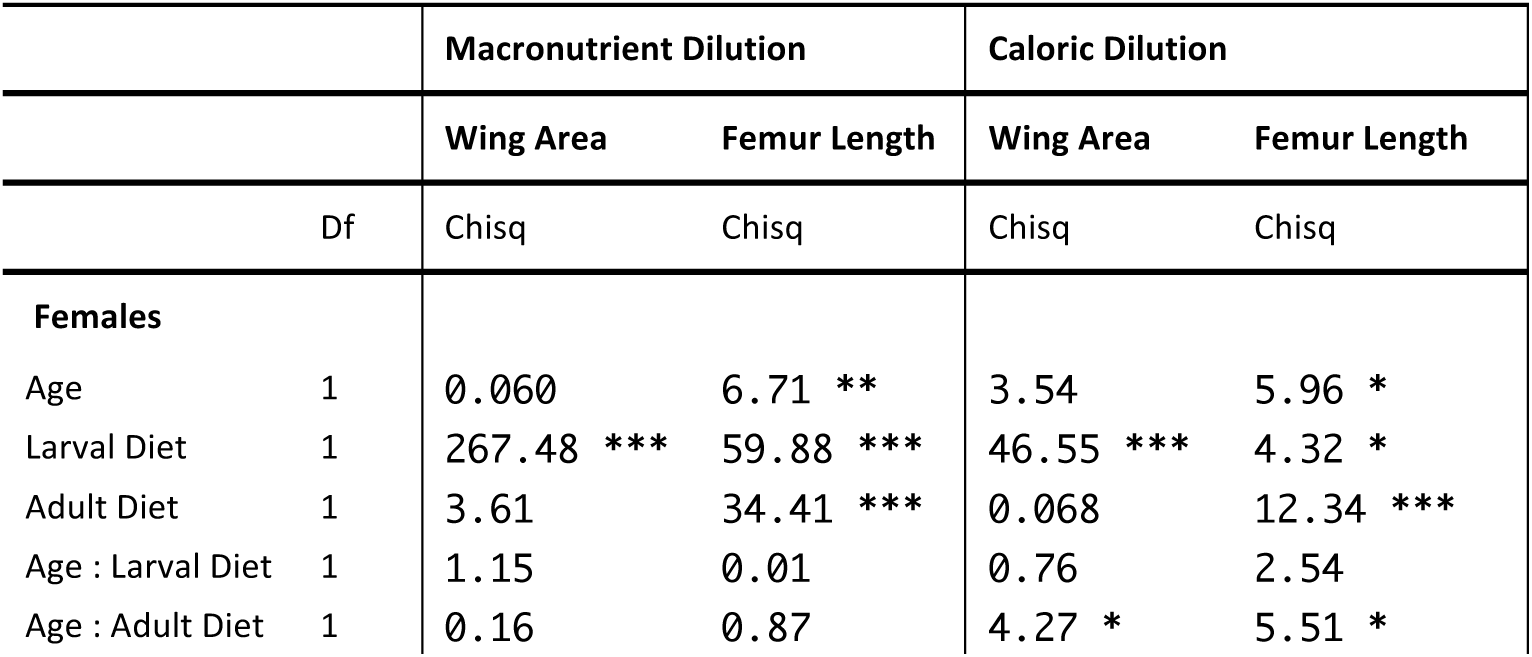
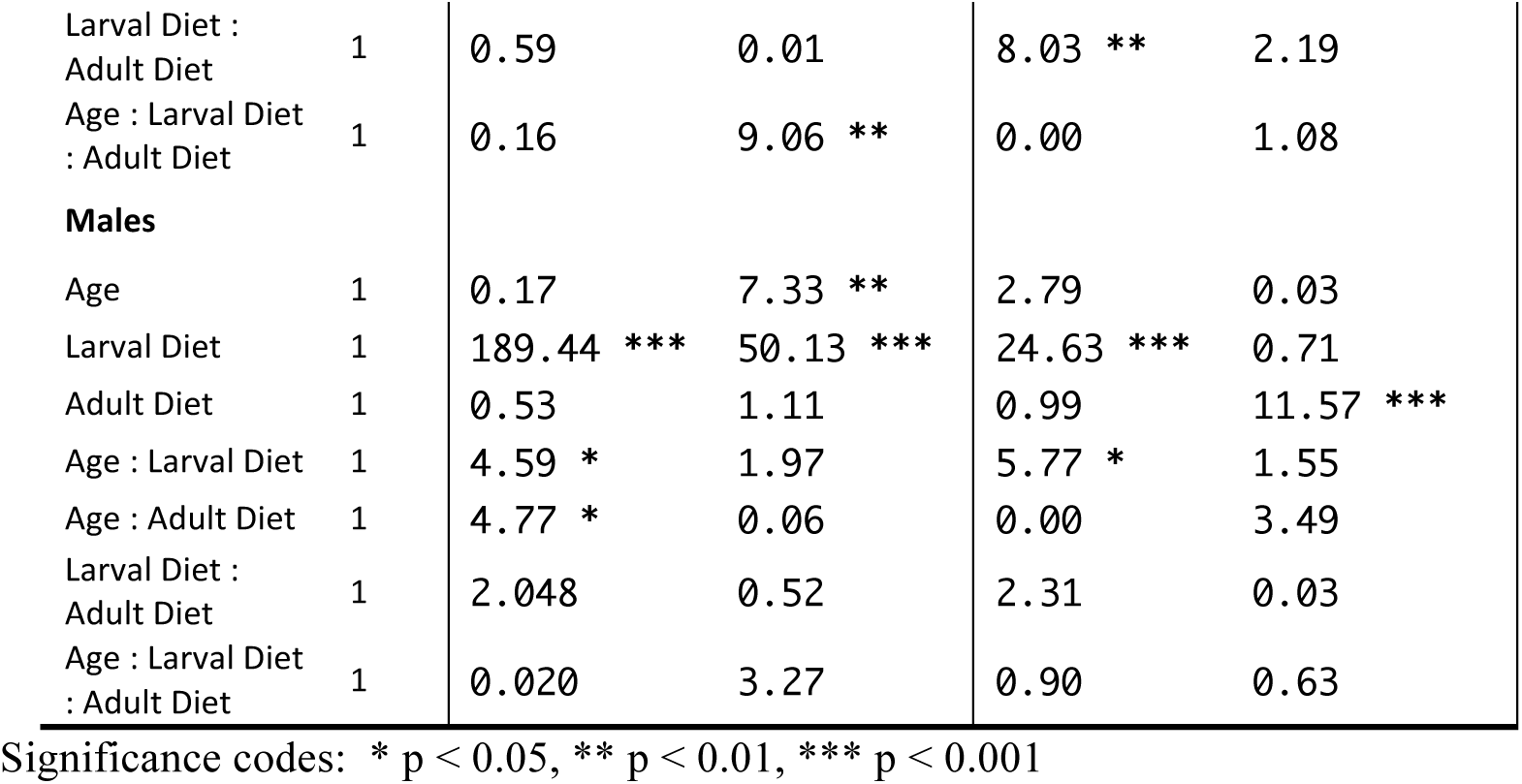
The effects of manipulating the larval and adult diets on wing area and femur length in females and males.

**Figure 4.**
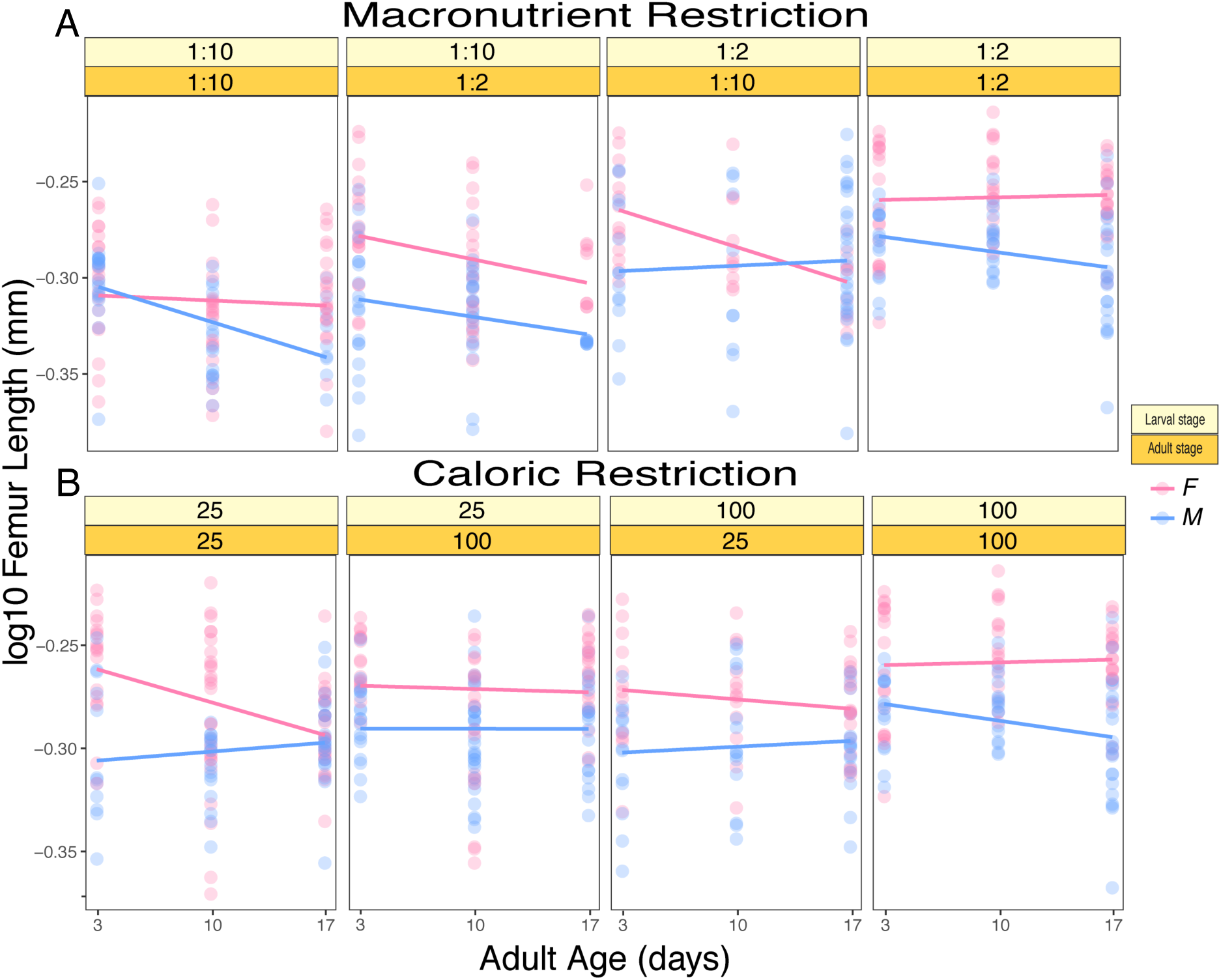
The length of the femur depends on the diet available to larvae and adults, and changes with age. A) shows the effects of the macronutrient composition of the larval and adult diet on femur length. B) shows the effects of the caloric composition of the larval and adult diet on femur length The stage at which treatments were imposed are indicated in the yellow-toned boxes: the larval nutrition in light yellow and the adult nutrition in darker yellow. The blue dots and lines represent the data for the adult males and the pink dots and lines represent the data for the adult females.

Female femur size further depended on age and the P:C ratio of the adult diet. As adults aged, their femurs became smaller (Figure 4A). The degree to which they decreased in size depended on both the diets on which they were reared as larvae, and the adult diet (Figure 4A, Table 5).

The P:C ratio of the larval diet also significantly affected male appendage size (Figure 3A, Table 5). Both the wings and femurs of males reared on the higher protein diet were larger than those from males reared on the low P:C diet. Similar to females, male femur length decreased with age (Figure 3A & 4A, Table 5). For males, we did not find significant interactions between age and diet for femur size. Male wing size did depend on interactions between age and the larval diet and age and the adult diet. The effects sizes were, however, very small.

#### 3.2.2. Caloric Manipulation

When we varied the caloric density of the diet most of the variance in wing size was explained by sex and the caloric content of the larval diet. Animals raised as larvae in the control diet (100%) eclosed with larger wings than those raised in the low-calorie diet (25%) (Figure 3B and Table 4). The effects of caloric restriction on wing size was smaller than those of macronutrient restriction (Figure 3A & 3B). Also, there were significant interactions between age and the caloric content of the larval diet, and caloric content of the larval and adult diets on adult wing size (Table 4).

Curiously, the caloric content of the larval diet did not significantly affect femur size. Instead, the caloric concentration of the adult diet and adult age contributed the most to variation in femur size (Figure 4B, Table 4). In addition, there were significant interactions between adult age and sex, the caloric content of the larval diet, age, and sex, and the caloric content of the adult diet, age, and sex. To unpack these complex interactions in both the wing and femur, we subdivided the data based on sex.

We identified significant interactions between the caloric content of the adult diet and age in both sexes on wing and femur size in females, and on wing size in males (Table 5). In females, femur size decreased with age, and in both sexes the adult diet significantly affected femur size where adults offered the higher calorie diets had slightly larger femurs (Figure 4B, Table 5). The degree to which femur size decreased with age in females depended on the caloric content of the adult diet (Table 5), with adult females on the low-calorie diets showing more pronounced reductions in femur size (Figure 4B).

Overall, the data from macronutrient and caloric manipulations demonstrate that the size of the wings is primarily affected by the larval diet, with little contribution of the adult diet and age to variation in wing size. In contrast, adult femur size changes more with age, and can depend on both the larval and adult diet. Surprisingly, this suggests that the size of adult appendages might not be as invariant as previously thought.

## 4. DISCUSSION

Many studies start from the principle that only larval diet matters for most measures of adult size. However, we know that body composition can change with adult diet (Skorupa et al., 2008; Verdú et al., 2010), suggesting that, depending on the measure of size, adult diet might also play important roles. We sought to understand the relative contribution of the larval versus adult diet to adult body weight and to the size of the adult appendages. Our hypothesis was that body weight would be affected by both larval and adult diet, while appendage size would be solely determined by larval diet.

Previous studies have also found that nutritional conditions during juvenile stages have the potential to affect life history traits in adult stages (Barrett et al., 2009; Dmitriew and Rowe, 2007; Metcalfe and Monaghan, 2001; Monaghan, 2008; Taborsky, 2006). This is particularly true of body size traits. For the most part, we found that the larval diet contributes more to differences in adult weight, wing area, and femur size in both males and females. These observations are consistent with previous studies in insects, which show that poor nutritional conditions in the larval stages affect the body composition and allometry of body components (Dmitriew and Rowe, 2007; Gotthard et al., 1994; Hahn, 2005; Nylin and Gotthard, 1998; Scharf et al., 2009; Scott et al., 2007; Stoks et al., 2006).

Our two types of dietary manipulation differed in the extent to which they changed adult size measures. By-in-large, manipulating the macronutrient content of the larval diet had larger effects on measures of adult size than manipulating the caloric content. Other studies have found that macronutrient content plays a greater role in shaping life history traits than caloric density. Both lifespan and fecundity in flies and other insects are more impacted by the P:C ratio of the diet than the caloric density (Carvalho et al., 2005; Fanson et al., 2012; Fanson and Taylor, 2012; Lee et al., 2008). The reason macronutrient balance is thought to matter for life history traits is that each trait requires specific amounts of either protein, carbohydrates, or both. For instance, the protein content of the larval diet appears to be the primary determinant of body size measures in several species of *Drosophila* (da Silva Soares et al., 2017; Kutz et al., 2019; Matavelli et al., 2015; Rodrigues et al., 2015; Shingleton et al., 2017). Taken together, it appears as though the source of calories, rather than the calories per se, shapes body size traits, with protein playing a central role in determining adult size.

However, the difference between dietary manipulations cannot be explained by protein concentration alone. The caloric content of the larval diet had more subtle effects on adult body size measures, even though the protein content of the low-calorie diet matched the protein content of the low P:C ratio diet. This suggests that the reduced size measures from the low P:C ratio diets arise due to the negative effects of sugar on growth. Other studies have proposed that sugar negatively impacts growth in the larval stages to reduce appendage size (Shingleton et al., 2017). This is likely because high levels of sugar in the diet impair a number of physiological functions, leading to reduced growth and delayed development (Havula et al., 2013).

While we found that low-calorie diets offered to larvae had more moderate effects on measures of body size, this is not to say that calories do not matter. In this study, we diluted calories to 25% the concentration of the standard diet for both the larvae and adults. Other studies have employed a broader range of nutritional compositions, using between 15-36 diets ranging in caloric content and macronutrient composition, to determine the effects on pharate adult weight and appendage size (Sentinella et al., 2013). These types of studies, which create a broad nutrient space, have tremendous power to uncover how traits are affected by dietary composition (Simpson and Raubenheimer, 2012). While this more extensive description of how size measures would be invaluable, implementing a fully factorial design that varies both the larval and the adult diet in this context would be challenging.

In addition to the effects of larval diet on adult size traits, we found that animals subjected to poor larval nutrition were able to increase their body weight if maintained on good quality diets during the adult stages. In females, some of these changes in body weight induced by good quality adult nutrition are likely to be due to increased ovarian mass, as well-fed females are known to produce higher numbers of eggs (Fanson and Taylor, 2012; Lee et al., 2008; Maklakov et al., 2008). Males also increased in mass with age, presumably due to changes in body composition such as altered body lipid, protein, and carbohydrate stores. These data support our hypothesis that size traits that are not constrained by the adult cuticle can respond more readily to the adult diet.

We also found that appendage size could change with time. Most notably, femur size decreased with age and changed with the composition of the adult diet in complex ways. The wing showed very little change with age or with adult diet, suggesting that these effects might be appendage specific. Further, unlike for body weight, rich adult diets were unable to increase femur size. This leads us to think that these small but significant changes in appendage size relate more to fly aging than to the effects of adult diet on appendage size. Interactions between diet and age may be a result of differences in the rate at which flies age under these differing conditions.

In conclusion, our study sought to uncover the relative contribution of the larval versus adult diet to measures of adult body size. We have found that while larval nutritional conditions play the dominant role in determining both adult body weight and appendage size in *D. melanogaster*, the adult diet can adjust body weight as the flies age. Unlike what has previously been found in humans and other animals (Barker and Thornburg, 2013; Barker et al., 1989; George et al., 2012; Leon et al., 1998; Painter et al., 2005), we found little evidence to support the idea that the nutritional history of the juvenile modifies the effects of the adult diet. Our study provides the basis for understanding how differences in diet across life stages induce metabolic changes to impact measures of adult body size, and for identifying stage-specific targets of diet’s action.

## 5. ACKNOWLEDGMENTS

We are grateful to the Mirth lab for their helpful discussions. This research was supported by funds from the Australian Research Council FT170100259 to CKM and the School of Biological Sciences, Monash University.

## 6. AUTHORS’ CONTRIBUTIONS

GP and CKM designed the experiments. GP and AC collected the data. GP and CKM completed the data analysis and wrote the manuscript.

## REFERENCES

Araújo, M.d.-S., Gil, L.H.S., e-Silva, A.d.-A., 2012. Larval food quantity affects development time, survival and adult biological traits that influence the vectorial capacity of Anopheles darlingi under laboratory conditions. Malaria Journal 11, 261.

Awmack, C.S., Leather, S.R., 2002. Host Plant Quality and Fecundity in Herbivorous Insects. Annual Review of Entomology 47, 817–844.

Bakker, K., 1959. Feeding period, growth, and pupation in larvae of *Drosophila melanogaster*. Entomologia Experimentalis et Applicata 2, 171–186.

Barker, D.J., Thornburg, K.L., 2013. The obstetric origins of health for a lifetime. Clinical Obstetrics and Gynecology 56, 511–519.

Barker, D.J., Winter, P.D., Osmond, C., Margetts, B., Simmonds, S.J., 1989. Weight in infancy and death from ischaemic heart disease. Lancet 2, 577–580.

Barrett, E.L., Hunt, J., Moore, A.J., Moore, P.J., 2009. Separate and combined effects of nutrition during juvenile and sexual development on female life-history trajectories: the thrifty phenotype in a cockroach. Proceedings of the Royal Society of London Series B 276, 3257–3264.

Behmer, S.T., 2009. Insect herbivore nutrient regulation. Annual Review of Entomology 54, 165–187.

Blackmore, M.S., Lord, C.C., 2000. The relationship between size and fecundity in *Aedes albopictus*. Journal of Vector Ecology 25, 212–217.

Boggs, C.L., Freeman, K.D., 2005. Larval Food Limitation in Butterflies: Effects on Adult Resource Allocation and Fitness. Oecologia 144, 353–361.

Bong, L.J., Neoh, K.B., Lee, C.Y., Jaal, Z., 2014. Effect of diet quality on survival and reproduction of adult *Paederus fuscipes* (Coleoptera:Staphylinidae). Journal of Medical Entomology 51, 752–759.

Callier, V., Nijhout, H.F., 2013. Body size determination in insects: a review and synthesis of size- and brain-dependent and independent mechanisms. Biological Reviews 88, 944–954.

Carvalho, G.B., Kapahi, P., Benzer, S., 2005. Compensatory ingestion upon dietary restriction in Drosophila melanogaster. Nature Methods 2, 813–815.

Chaudhury, M.F.B., 1989. Nutritional Ecology of Insects, Mites, Spiders, and Related Invertebrates. International Journal of Tropical Insect Science 10, 253–253.

Colasurdo, N., Gelinas, Y., Despland, E., 2009. Larval nutrition affects life history traits in a capital breeding moth. Journal of Experimental Biology 212, 1794–1800.

da Silva Soares, N.F., Alves, A.N., Beldade, P., Mirth, C.K., 2017. Adaptation to novel nutritional environments: larval performance, foraging decisions, and adult oviposition choices in *Drosophila suzukii*. BMC Ecology 17, 21.

David, J.R., 1970. Le nombre d’ovarioles chez *Drosophila melanogaster*: relation avec la fécondité et valeur adaptive. Archives de Zoologie Expérimentale et Générale 111, 357–370.

Dmitriew, C., Rowe, L., 2007. Effects of early resource limitation and compensatory growth on lifetime fitness in the ladybird beetle (Harmonia axyridis). Journal of Evolutionary Biology 20, 1298–1310.

Emlen, D.J., Nijhout, H.F., 1999. Hormonal control of male horn length dimorphism in the dung beetle *Onthophagus taurus* (Coleoptera: Scarabaeidae). Journal of Insect Physiology 45, 45–53.

Fanson, B.G., Fanson, K.V., Taylor, P.W., 2012. Cost of reproduction in the Queensland fruit fly: Y-model versus lethal protein hypothesis. Proceedings of the Royal Society of London Series B 279, 4893–4900.

Fanson, B.G., Taylor, P.W., 2012. Protein:carbohydrate ratios explain life span patterns found in Queensland fruit fly on diets varying in yeast:sugar ratios. Age (Dordr) 34, 1361–1368.

Galikova, M., Klepsatel, P., 2018. Obesity and aging in the *Drosophila* model. International Journal of Molecular Sciences 19, 1896.

George, L.A., Zhang, L., Tuersunjiang, N., Ma, Y., Long, N.M., Uthlaut, A.B., Smith, D.T., Nathanielsz, P.W., Ford, S.P., 2012. Early maternal undernutrition programs increased feed intake, altered glucose metabolism and insulin secretion, and liver function in aged female offspring. American Journal of Physiology-Regulatory, Integrative and Comparative Physiology 302, R795–804.

Gobbi, P., Martínez-Sánchez, A., Rojo, S., 2013. The effects of larval diet on adult life-history traits of the black soldier fly, Hermetia illucens (Diptera: Stratiomyidae). European Journal of Entomology 110, 461–468.

Gotthard, K., Nylin, S., Wiklund, C., 1994. Adaptive variation in growth rate: life history costs and consequences in the speckled wood butterfly, Pararge aegeria. Oecologia 99, 281–289.

Grangeteau, C., Yahou, F., Everaerts, C., Dupont, S., Farine, J.P., Beney, L., Ferveur, J.F., 2018. Yeast quality in juvenile diet affects *Drosophila melanogaster* adult life traits. Scientific Reports 8, 13070.

Hahn, D.A., 2005. Larval nutrition affects lipid storage and growth, but not protein or carbohydrate storage in newly eclosed adults of the grasshopper *Schistocerca americana*. Journal of Insect Physiology 51, 1210–1219.

Havula, E., Teesalu, M., Hyötyläinen, T., Seppälä, H., Hasygar, K., Auvinen, P., Oresic, M., Sandmann, T., Hietakangas, 2013. Mondo/ChREBO-Mlx-regulated transcriptional network is essential for dietary sugar tolerance in *Drosophila*. PLoS Genetics 9, e1003438.

Helm, B.R., Rinehart, J.P., Yocum, G.D., Greenlee, K.J., Bowsher, J.H., 2017. Metamorphosis is induced by food absence rather than a critical weight in the solitary bee, *Osmia lignaria*. Proceedings of the National Academy of Science USA 114, 10924–10929.

Jensen, K., Mayntz, D., Toft, S., Clissold, F., Hunt, J., Raubenheimer, D., Simpson, S.J., 2012. Optimal foraging for specific nutrients in predatory beetles. Proceedings of the Royal Society of London Series B 279, 2212–2218.

Jones, S.A., Raubenheimer, D., 2001. Nutritional regulation in nymphs of the German cockroach, *Blattella germanica*. Journal of Insect Physiology 47, 1169–1180.

Kutz, T.C., Sgrò, C.M., Mirth, C.K., 2019. Interacting with change: diet mediates how larvae respond to their thermal environment. Functional Ecology 33, 1940–1951.

Lee, K.P., Simpson, S.J., Clissold, F.J., Brooks, R., Ballard, J.W., Taylor, P.W., Soran, N., Raubenheimer, D., 2008. Lifespan and reproduction in Drosophila: New insights from nutritional geometry. Proceedings of the National Academy of Science U S A 105, 2498–2503.

Leon, D.A., Lithell, H.O., Vagero, D., Koupilova, I., Mohsen, R., Berglund, L., Lithell, U.B., McKeigue, P.M., 1998. Reduced fetal growth rate and increased risk of death from ischaemic heart disease: cohort study of 15 000 Swedish men and women born 1915-29. BMJ 317, 241–245.

Magnarelli, L.A., Leprince, D.J., Burger, J.F., Butler, J.F., 1982. Oviposition Behavior and Fecundity in Chrysops Cincticornis (Diptera: Tabanidae)1. Journal of Medical Entomology 19, 597–600.

Maklakov, A.A., Simpson, S.J., Zajitschek, F., Hall, M.D., Dessmann, J., Clissold, F., Raubenheimer, D., Bonduriansky, R., Brooks, R.C., 2008. Sex-specific fitness effects of nutrient intake on reproduction and lifespan. Current Biology 18, 1062–1066.

Matavelli, C., Carvalho, M.J.A., Martins, N.E., Mirth, C.K., 2015. Differences in larval nutritional requirements and female oviposition preference reflect the order of fruit colonization of *Zaprionus indianus* and *Drosophila simulans*. Journal of Insect Physiology 82, 66–74.

Mayntz, D., Raubenheimer, D., Salomon, M., Toft, S., Simpson, S.J., 2005. Nutrient-specific foraging in invertebrate predators. Science 307, 111–113.

Metcalfe, N.B., Monaghan, P., 2001. Compensation for a bad start: grow now, pay later? Trends in Ecology and Evolution 16, 254–260.

Monaghan, P., 2008. Early growth conditions, phenotypic development and environmental change. Philosophical Transactions of the Royal Society of London Series B: Biological Sciences 363, 1635–1645.

Nijhout, H.F., 2003. The control of body size in insects. Developmental Biology 261, 1–9.

Nijhout, H.F., Grunert, L.W., 2010. The cellular and physiological mechanism of wing-body scaling in Manduca sexta. Science 330, 1693–1695.

Nijhout, H.J., Riddiford, L.M., Mirth, C.K., Shingleton, A.W., Suzuki, Y., Callier, V., 2014. The developmental control of size in insects. WIREs Developmental Biology 3, 113–134.

Nylin, S., Gotthard, K., 1998. Plasticity in life-history traits. Annu Rev Entomol 43, 63–83.

Painter, R.C., Roseboom, T.J., Bleker, O.P., 2005. Prenatal exposure to the Dutch famine and disease in later life: an overview. Reproductive Toxicology 20, 345–352.

Rodrigues, M.A., Martins, N.E., Balancé, L.F., Broom, L.N., Dias, A.J.S., Fernandes, A.S.D., Rodrigues, F., Sucena, É., Mirth, C.K., 2015. *Drosophila melanogaster* larvae make foraging choices to minimize developmental time. Journal of Insect Physiology 81, 69–80.

Scharf, I., Filin, I., Ovadia, O., 2009. A trade-off between growth and starvation endurance in a pit-building antlion. Oecologia 160, 453–460.

Scott, D.E., Casey, E.D., Donovan, M.F., Lynch, T.K., 2007. Amphibian lipid levels at metamorphosis correlate to post-metamorphic terrestrial survival. Oecologia 153, 521–532.

Sentinella, A.T., Crean, A.J., Bonduriansky, R., 2013. Dietary protein mediates a trade-off between larval survival and the development of male secondary sexual traits. Functional Ecology 27, 1134–1144.

Shafiei, M., Moczek, A.P., Nijhout, H.F., 2001. Food availability controls the onset of metamorphosis in the dung beetle *Onthophagus taurus* (Coleoptera: Scarabaeidae). Physiological Entomology 26, 173–180.

Shingleton, A.W., Masandika, J., Thorsen, L., Zhu, Y., Mirth, C.K., 2017. The sex-specific effects of diet quality versus quantity on size and shape in *Drosophila melanogaster*. Royal Society Open Science 4, 170375.

Simpson, S.J., James, S., Simmonds, M.S., Blaney, W.M., 1991. Variation in chemosensitivity and the control of dietary selection behaviour in the locust. Appetite 17, 141–154.

Simpson, S.J., Raubenheimer, D., 2012. The nature of nutrition: A unifying framework from animal adaptation to human obesity. Princeton University Press, Princeton.

Skorupa, D.A., Dervisefendic, A., Zwiener, J., Pletcher, S.D., 2008. Dietary composition specifies consumption, obesity, and lifespan in Drosophila melanogaster. Aging Cell 7, 478–490.

Souza, R.S., Virginio, F., Riback, T.I.S., Suesdek, L., Barufi, J.B., Genta, F.A., 2019. Microorganism-Based Larval Diets Affect Mosquito Development, Size and Nutritional Reserves in the Yellow Fever Mosquito Aedes aegypti (Diptera: Culicidae). Frontiers in Physiology 10.

Stoks, R., De Block, M., McPeek, M.A., 2006. Physiological costs of compensatory growth in a damselfly. Ecology 87, 1566–1574.

Taborsky, B., 2006. The influence of juvenile and adult environments on life-history trajectories. Proceedings of the Royal Society of London Series B 273, 741–750.

Tomberlin, J.K., Sheppard, D.C., Joyce, J.A., 2002. Selected Life-History Traits of Black Soldier Flies (Diptera: Stratiomyidae) Reared on Three Artificial Diets. Annals of the Entomological Society of America 95, 379-386, 378.

Verdú, J.R., Casa, J.L., Lobo, J.M., Numa, C., 2010. Dung beetles eat acorns to increase their ovarian development and thermal tolerance. PLoS One 5, e10114.

